# High Fat Diet Blunts Stress-Induced Hypophagia and Activation of Glp1r Dorsal Lateral Septum Neurons in Male but not in Female Mice

**DOI:** 10.1101/2022.02.08.479553

**Authors:** Michelle B. Bales, Samuel W. Centanni, Joseph R. Luchsinger, Kaitlyn Ginika Nwaba, Isabella M. Paldrmic, Danny G. Winder, Julio E. Ayala

## Abstract

While stress typically reduces caloric intake (hypophagia) in chow-fed rodents, presentation of palatable, high calorie substances during stress can increase caloric consumption (i.e. “comfort feeding”) and promote obesity. However, little is known about how obesity itself affects feeding behavior in response to stress and the mechanisms that can influence stress-associated feeding in the context of obesity. We show that lean male mice display the expected hypophagic response following acute restraint stress, but obese male mice are resistant to this acute stress-induced hypophagia. Activation of the glucagon-like peptide-1 (Glp1) receptor (Glp1r) in various brain regions leads to hypophagia in response to stress. Here we show that Glp1r-positive neurons in the dorsal lateral septum (dLS) are robustly activated during acute restraint stress in lean but not in obese male mice. This raises the possibility that activation of dLS Glp1r neurons during restraint stress contributes to subsequent hypophagia. Supporting this, we show that chemogenetic inhibition of dLS Glp1r neurons attenuates acute restraint stress hypophagia in male mice. Surprisingly, we show that both lean and obese female mice are resistant to acute restraint stress-induced hypophagia and activation of dLS Glp1r neurons. Taken together, these results suggest that dLS Glp1r neurons contribute to the hypophagic response to acute restraint stress in male mice, but not in female mice, and that obesity disrupts this response in male mice. Broadly, these findings show sexually dimorphic mechanisms and feeding behaviors in lean vs. obese mice in response to acute stress.

## 1.0 Introduction

Stress is a ubiquitous environmental factor that can affect feeding behavior. Although stress reduces caloric intake and promotes weight loss in some individuals, the typical response to stress is overconsumption and weight gain (Oliver & Wardle, 1999; Weinstein et al., 1997). Research on the relationship between stress and obesity has primarily focused on how stress contributes to the development of obesity. Much less is known about how obesity itself affects the response to stress. Clinical studies suggest that obese individuals are more likely to increase caloric consumption and engage in “emotional eating” as a means to cope with stress (Boggian et al., 2015). Elucidating mechanisms that regulate caloric intake in response to stress in lean and obese states could identify interventions to reduce overconsumption during times of stress.

Glucagon-like peptide-1 (Glp1) has emerged as a key molecule that regulates the response to stress. Although it is widely recognized as a gut secreted peptide that stimulates insulin secretion by engaging a pancreatic β-cell Glp1 receptor (Glp1r), Glp1 is also secreted from preproglucagon (PPG) neurons in the nucleus of the tractus solitarius (NTS; Holt et al., 2019). Activation of NTS PPG neurons reduces food intake and body weight in rodents (Gaykema et al., 2017; Holt et al., 2019). This is likely due, at least in part, to subsequent stimulation of Glp1r signaling in brain regions innervated by NTS PPG neurons, such as the lateral parabrachial nucleus (LpBN) and hypothalamus, since targeting Glp1r agonists to these brain regions also reduces food intake and body weight (Alhadeff et al., 2014; López-Ferreras et al., 2018). Reduced food intake (hypophagia) is a typical stress response in rodents, and several lines of evidence suggest that endogenous Glp1 action in the brain plays an important role in this behavior. Central administration of Glp1 induces c-fos expression, a marker for neuronal activity, in stress-related corticotropin releasing hormone (CRH)-producing neurons of the paraventricular nucleus (PVN; Larsen et al., 1997) and stimulates the hypothalamic-pituitary-adrenal (HPA) axis in rats (Kinzig et al., 2003). Acute stress causes activation of PPG neurons in the NTS in rats (Maniscalco et al., 2015) and mice (Terrill et al., 2019), and chemogenetic inhibition of NTS PPG neurons attenuates acute stress-induced hypophagia (Holt et al., 2019). Additionally, intracerebroventricular (ICV) delivery of the Glp1r antagonist Exendin (9-39) (Ex9) reduces the ability of acute restraint stress to promote hypophagia (Maniscalco et al., 2015). Targeting Ex9 to specific brain regions such as the bed nucleus of the stria terminalis (BNST; Ghosal et al., 2017) and the PVN (Zheng et al., 2019) shows that inhibition of the Glp1r in these regions attenuates the hypophagic response to stress.

The LS is another brain region that may be targeted by Glp1 to regulate feeding behavior in response to stress. The LS is a relay center for various brain regions, and it plays a key role in modulating behavioral responses to anxiety and stress (Olds & Milner, 1954; Singewald et al., 2011). Restraint stress increases c-fos expression in the LS in mice (Patel, 2005; Anthony et al., 2014; Mitra et al., 2014), and direct activation of Glp1r-expressing neurons in the LS reduces food intake (Azevedo et al., 2020). This suggests that LS Glp1 action promotes hypophagia in response to restraint stress. This is further supported by studies showing that inhibition of LS Glp1r signaling, specifically in the dorsal LS (dLS), attenuates restraint stress-induced hypophagia in rodents (Terrill et al., 2018; Terrill et al, 2019). Interestingly, restraint stress hypophagia is attenuated in rats given intermittent access to sucrose prior to stress, and this is associated with blunted activation of LS neurons compared to chow-fed controls (Martin & Timofeeva, 2010; Michel et al., 2003). Taken together, these results suggest that LS Glp1r neurons promote restraint stress hypophagia, and the activation of these neurons in response to restraint stress is attenuated by exposure to palatable, high calorie substances.

The present studies assessed the effect of obesity on food intake and dLS Glp1r neuronal activity in response to acute restraint stress in mice and whether dLS Glp1r neurons are necessary for stress-induced hypophagia. Our findings demonstrate that obese male mice are resistant to acute restraint stress-induced hypophagia, and this is associated with decreased activation of dLS Glp1r neurons. Furthermore, inhibition of dLS Glp1r neuronal activity blunts acute restraint stress-induced hypophagia in lean male mice, suggesting that dLS Glp1r neurons mediate hypophagia following acute restraint stress. Interestingly, we find that both lean and obese female mice are resistant to the hypophagic response following acute restraint stress, and they are also resistant to the activation of dLS Glp1r neurons during restraint.

## 2. Materials and methods

### 2.1 Subjects

All mice were housed in the Vanderbilt University Medical Center animal vivarium and kept on a standard 12hr/12hr light/dark cycle. Mice had *ad libitum* access to distilled water and were either maintained on a chow diet (LabDiet, 5L0D) from the time of weaning or were given a high fat diet (HFD, Research Diets Inc., D12492) from six weeks of age. All animals were 4 to 6 months old at the time of testing. Experimental procedures were approved by The Institutional Animal Care and Use Committee at Vanderbilt University.

### 2.2 Handling Acclimation

To prevent stress from handling alone, mice underwent a handling acclimation procedure, modified from Olsen & Winder (2010), for five days prior to testing. On Day 1, the experimenter placed a newly-gloved hand in the cage for 90 s and the animal was allowed to explore the gloved hand. This only occurred on Day 1. The mouse was then placed on top of the hand which gently lifted a few inches from the cage floor for a few seconds and was then placed back down slowly for the mouse to dismount the hand. This movement was repeated 5-10 times depending on the demeanor of the mouse (e.g. if the mouse was slow to move away from the hand, indicating acclimation to the hand, the experimenter would repeat this only 5 times, but if the mouse was quick to dismount the hand, the experimenter would lift the mouse 10 times to acclimate further). On the final hand lift, the mouse was held higher in the air (∼20 cm) for 5 s on Day 1, 10 s on Day 2, and 15 s on Day 3. On the last two days of the handling acclimation procedure, the mouse was simply placed into the hand, stroked a few times, and then weighed.

### 2.3 Food Intake Following Restraint Stress

Chow- and HFD-fed male and female C57BL/6J mice (#000664, Jackson Laboratories) were individually housed at least one week before being singly-housed in a Promethion metabolic system (Sable Systems, Inc.) for at least one week. Following the handling acclimation procedure (see 2.2) mice underwent restraint stress for one hour prior to dark onset. Chow-fed mice were gently guided into a 50 mL conical tube (1 inch diameter) that was modified to have the same number of air holes and tail/nose holes as restrainers for obese mice (at least 1.5 inches in diameter; Plas Labs, #552-BSRR). Control (no restraint stress) mice were picked up by the tail and placed back down in the cage. Food intake, water intake, energy expenditure, substrate oxidation, and locomotor activity were recorded continuously every 5 min overnight. Mice that spilled or cashed >0.5 g of food in a 5 min bin were removed from the analysis.

### 2.4 LS Glp1r Neuronal Activity During Restraint Stress

Real-time activity of dLS Glp1r neurons during restraint stress was measured using fiber photometry. Struggle behavior was assessed during the fiber photometry sessions using DeepLabCut (DLC) video analysis as previously described (Luchsinger et al., 2021).

#### 2.4.1 Surgery

Chow- and HFD-fed male and female mice expressing Cre recombinase under the control of the *glp1r* locus (Glp1r-Cre; Williams et al., 2016) were deeply anesthetized and secured into an Angle Two stereotaxic instrument (Leica Biosystems). A craniotomy was performed under isoflurane anesthesia (2-3%). The Cre-dependent intracellular Ca2+ sensor (GCaMP7f) or control GFP (see Table 1) was then infused into the dLS (300 nL in the left hemisphere; 50 nL/second) using a microinjection syringe pump and a 33G needle (WPI) at a 22.05° angle. The coordinates for the injection were +0.5 mm rostral, +/-0.3 mm lateral, and -3.0 mm ventral. A borosilicate mono fiber-optic cannula (Doric Lenses Inc.) with a 400 µm core diameter was then implanted into the dLS using the same coordinates as the GCaMP7 and was secured to the skull with two small screws (Plastics One) and dental cement. Mice were given three weeks to recover and to allow time for the GCaMP7 to express before testing.

**Table 1.**
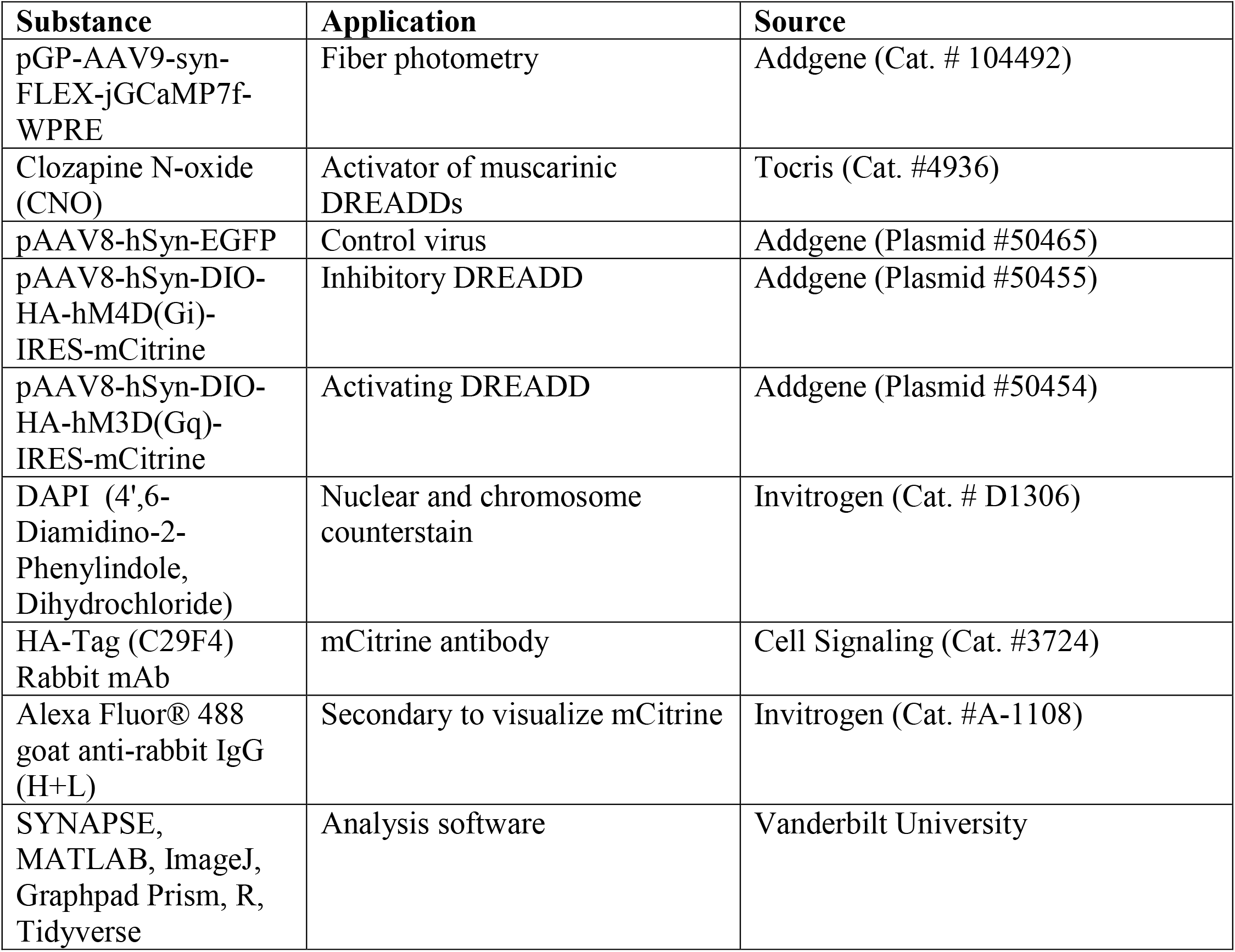
Materials used

#### 2.4.2 Recording Procedure

Mice underwent the handling acclimation protocol (see 2.2). Mice were also acclimated to the fiber-optic patch cord, which attaches to the fiber-optic cannula, two days prior to testing. On the test day, animals acclimated to the testing room for at least one hour. Each mouse was connected to the fiber-optic patch cord and allowed five minutes of acclimation before recording began. The recordings were captured with an LED-based fiber photometry system (Tucker-Davis Technologies, TDT, Model is RZ5P) with Synapse acquisition and analysis software that uses real-time lock-in amplification and signal demodulation to detect in vivo fluorescence lifetime measurements and photometry recordings through optical fibers. Each recording consisted of a single-mode excitation fiber originating from a 470nM wavelength LED, and the multimode detection fiber that was surgically implanted and collects bulk fluorescence induced by the GCaMP7. Recordings were made during the restraint period as well as 10 minutes before and 10 minutes after the restraint stress with no pause in recording. Mice were also video recorded (Logitech PRO HD Webcam C920) for assessment of struggle behavior (see 2.4.5 below). Restrainers varied in diameter based on the weight of the mouse to help account for variability in level of restraint. Mice between 21 and 30.9 g were gently guided into a clear acrylic restraint device described in Luchsinger et al. (2021) with a 1 in. diameter (Vanderbilt Kennedy Center Scientific Instrumentation Core). Mice that were 31 to 40.9 g went into a restrainer that was at least 1.5 inches in diameter (Plas Labs, #552-BSRR).

#### 2.4.3 Fiber-Optic Cannula and GCaMP7 confirmation

After testing was complete, animals were deeply anesthetized and transcardially perfused with PBS followed by 4% paraformaldehyde (PFA). Brains were placed in 4% PFA for at least 24 hours then transferred to a 20% sucrose in PBS solution to cryoprotect the brains for at least 48 hours, all at 4°C. Brains were carefully blocked and then mounted on a freezing microtome and sectioned coronally at 35 µm. Sections were then mounted on Superfrost slides and coverslipped with ProLong Gold Antifade Mountant in order to histologically confirm the placement of the GCaMP7 and the fiber-optic cannula. Sections were examined under a fluorescent microscope for correct cannula placement and fluorescent signal of GCaMP7. Only animals with correct fiber-optic cannula placement were included in the analysis.

#### 2.4.4 Fiber Photometry Data Analysis

Each recording was broken down into seven segments comprising of 8-minutes of recording, leaving two minutes to account for time it took to get the mouse into the restrainer, and then analyzed separately to account for a natural drift in baseline. Raw photon count was converted to ΔF/F_0_ using a segmented normalization procedure with a bin of 4 s and Ca2+ transients was determined using a Savitsky–Golay filter and empirically identified kinetic parameters using the MLspike algorithm in MATLAB (Harris et al., 2018). The mLspike algorithm, which is a validated way to impute spike frequency based on Ca2+ transients (Deneux et al., 2016), was used to measure spike frequency. Area under the curve was measured and statistics were calculated in Prism 9 (GraphPad). Group differences were assessed with two-way ANOVAs and Bonferroni’s multiple comparisons test. P values of < .05 were considered significant.

#### 2.4.5 Head Movement Analysis

Videos from fiber photometry test day were cropped to include only the time when the animal was in the restrainer. One minute was taken off the beginning and the end of each video to account for some variation in start and stop times across recordings. These 58 min videos were then analyzed for head movements using DLC as described in Luchsinger et al. (2021), and briefly described here. Tail movement analysis was omitted due to the restrainers for larger mice obscuring the tails from view. Regardless, head movement accounts for the majority of a struggle behavior (Luchsinger et al., 2021). Following hand-scoring of head movements on similarly restrained mice, where bouts were separated by <0.7 s pause, DLC was used to track the same head movements. DLC allows for automated video scoring by tracking points across time using similar methods for scoring manually (Nath et al., 2017; Mathis et al., 2018). The point that was used to track head movements, and was consistent across recordings, was a small piece of green tape attached to the fiber-optic patch cable. DLC was trained to automatically track this point after manually locating the point on at least 495 images. The statistical software R (R Foundation for Statistical Computing, 2019) and the “tidyverse” package (Wickham et al., 2019) was used to make the point (the green tape) an X/Y position based on pixels into a speed of the head movement for each frame of the video (10 fps). Head movements were counted when the frame had a speed greater than one standard deviation above the lower 95% of frame speeds. Normalizing the speed of the head movement in this way allowed for more consistency across animals to account for small differences in restrainer and camera placement, though the experimenter took care to mitigate changes in camera and restrainer placement during testing. AUC and maximum peak amplitude were calculated for the 5 s after bout onset to normalize across bouts. Maximum peak was time-locked to the fiber photometry recording in MATLAB. Head movement figures were created in R with the ggplot2 package (Wickham, 2016).

### 2.5 Chemogenetic Modulation of dLS Glp1r Neurons

In Glp1r Cre+ male mice, activating (pAAV8-hSyn-DIO-HA-hM3D(Gq)-IRES-mCitrine; Addgene) or inhibiting (pAAV8-hSyn-DIO-HA-hM4D(Gi)-IRES-mCitrine; Addgene) designer receptors exclusively activated by designer drugs (DREADD) or control (pAAV8-hSyn-EGFP; Addgene) were delivered into the dLS as described for GCaMP7 (see 2.4.1) except that 300 nL of DREADD/control AAV was injected into each hemisphere, totaling 600 nL (50 nL/sec) via a microinjection syringe pump and a 33G needle (WPI) at a 22.05° angle. Surgery was performed under anesthesia (2-3% isoflurane), and mice were given at least one week to recover and to allow time for the DREADD to express before testing.

Following recovery, mice were individually housed at least one week before going into the Promethion and then acclimated to the Promethion for one week and underwent the handling acclimation procedure (see 2.2). Mice also acclimated to injections with saline (i.p.) four days prior to test days. Treatment groups were balanced based on bodyweight. Activating DREADD experiments had a crossover design where the same animal was injected with saline on Test Day 1 and the DREADD ligand clozapine N-oxide (CNO, 2 mg/kg) on Test Day 2 or vice versa with four days between Test Days. Inhibiting DREADD experiments were also performed with a crossover design in which mice were restrained with either saline or CNO injections on Test Days 1 and 4 or remained unrestrained with saline or CNO injections on Test Days 2 and 3.

Saline and CNO were injected 30 minutes before restraint stress or no restraint stress which was administered one hour before dark onset for Test Days 1-3. There were two days between test days during which all mice continued receiving i.p. saline injections. Test Day 4 was followed by transcardial perfusion, so no food intake was measured. Only mice with confirmed DREADD expression in dLS were included in the analyses and group differences were assessed with two-way ANOVAs and Bonferroni’s multiple comparisons test. P values of < .05 were considered significant. Outlier tests were conducted by measuring mean food intake every 2 h during the 12h dark cycle (6 time points) Mice with food intake values greater than 2x standard deviation from the mean in 3 or more consecutive time points were excluded as outliers.

#### 2.5.3 DREADD Confirmation

After food intake testing was complete, animals were anesthetized and transcardially perfused with PBS followed by freshly prepared 4% paraformaldehyde (PFA). Brains were then extracted and placed in 4% PFA for at least 24 hrs then transferred to a cryoprotectant solution of 20% sucrose in PBS for at least 48 hrs, all at 4°C. Brains were then coronally sectioned on a freezing microtome at 35 µm.

Sections were selected at 25, 50, and 75% the length of dLS, at approximately 1.0, 0.5, and 0.0 mm AP based on anatomical landmarks (Franklin and Paxinos, 2008) for free-floating immunohistochemistry. To stain for the HA-Tag to visualize the DREADD expression, sections first permeabilized in 0.4% Triton X-100 in PBS for one hour at room temperature. Sections then sat in blocking buffer (1% Bovine Serum Albumin, 5% Normal Goat Serum [NGS] in the same Triton X-100 solution) for an hour at room temperature before sitting in the primary HA-Tag rabbit antibody (1:500) in blocking buffer for 48 hours at 4°C. Following rinses (3 × 10 min) in the Triton X-100 solution, sections were kept in the goat anti-rabbit Alexa Fluor 488 secondary (1:250) in blocking buffer solution for two hours at room temperature. Rinses (3 × 10 min) in the Triton X-100 solution occurred before applying a DAPI (1:25000) in PBS solution (15 min). One more rinse, this time in PBS (10 min), was administered before sections were mounted onto glass slides and allowed to dry overnight. Slides were then coverslipped with ProLong Gold Antifade Mountant. Slides were imaged on the LSM710 confocal at 5x to confirm DREADD placement.

## 3.0 Results

### 3.1 Energy Balance Parameters in Lean and Obese Mice Following Restraint Stress

After twelve weeks on HFD male mice were obese compared to chow-fed males (38.7±6.3g vs. 29.1± 3.0g). Female mice were also obese following HFD feeding compared to female chow-fed mice (32.6±6.1g vs. 23.3±2.9g). As expected, food intake decreased following restraint stress in lean male mice shown by a main effect of the one hour restraint on chow intake over the twelve hours following dark onset (Two-way ANOVA: F [144, 2755] = 1.759; P < 0.0001; Figure 1A). Consistent with previous studies (Terrill et al., 2019), decreased food intake was not evident until ∼4h post restraint stress. Indeed, when food intake is plotted every 2 hours, t-tests with Bonferroni correction revealed significant effects at 10 and 12 hours post dark onset (P = 0.0098 at 10 hours, P = 0.0112 at 12 hours; Figure 1A inset). Contrasting the phenotype in lean mice, there was no effect of the one hour restraint on food intake in obese male mice over the same twelve hours following dark onset (Two-way ANOVA: F [144, 2755] = 0.08609; P > 0.9999; Figure 1B). Interestingly, there was no effect of the restraint stress on food intake in female mice whether they were lean (Two-way ANOVA: F [144, 2030] = 0.3234; P > 0.9999; Figure 1C) or obese (Two-way ANOVA: F [144, 2030] = 0.1932; P > 0.9999; Figure 1D). These results show that obese male mice are resistant to acute restraint stress-induced hypophagia, but female mice do not display acute restraint stress-induced hypophagia regardless of metabolic status.

**Figure 1.**
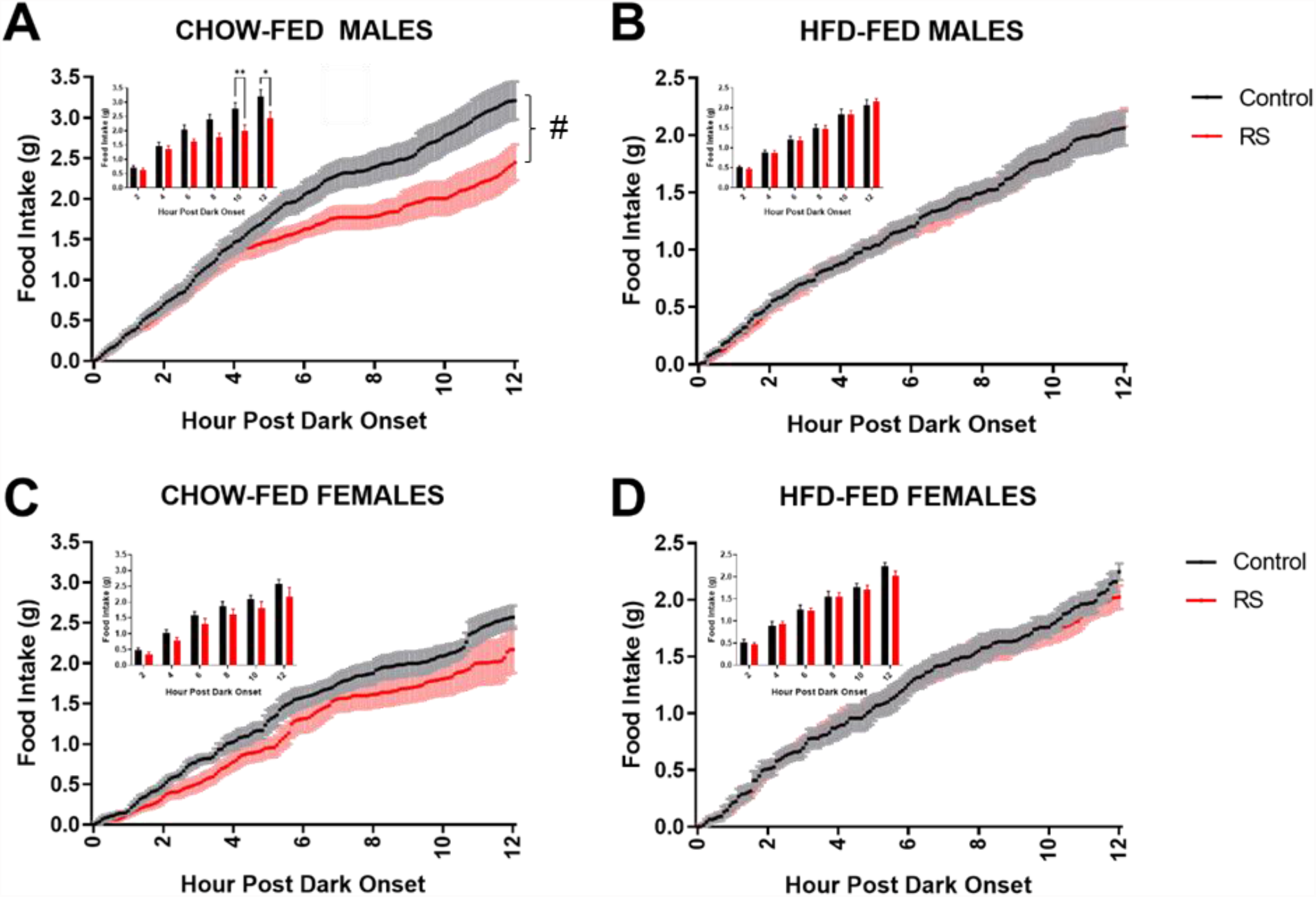
Restraint Stress Hypophagia in Lean Male Mice but Not in Obese Male or Lean or Obese Female Mice. Food intake following no restraint (Control; black lines/bars) or restraint stress (RS; red lines/bars) which occurred for one hour prior to dark onset. Male (A, B) and female (C, D) wild-type mice were either chow-fed (A, C) or HFD-fed (B, D). #P < 0.0001 in a Two-way ANOVA; *P < 0.05 & **P < 0.01 in a t-test. N = 9-12/group.

Restraint stress significantly increases energy expenditure (EE) over 12 hours in lean male mice (Two-way ANOVA: F (144, 3190) = 1.362, P = 0.0032; Figure 2A) and tended to increase EE in obese male mice (Two-way ANOVA: F (144, 2755) = 1.164, P = 0.0935; Figure 2B). There was a significant increase in EE during the one hour restraint stress period for both lean (unpaired t-test, P < 0.0001; Figure inset 2A) and obese male mice (unpaired t-test, P = 0.0092; Figure inset 2B). Elevated EE during restraint could be due to increased breathing and muscle contractions as the mice struggle within the restrainer. The observation that EE is increased during restraint stress in both lean and obese male mice suggests that both lean and obese mice experience stress equally, and the absence of stress-induced hypophagia in obese mice is not due to a difference in the stress experience. A more direct measure of stress (i.e. struggle behavior) will be discussed below.

**Figure 2.**
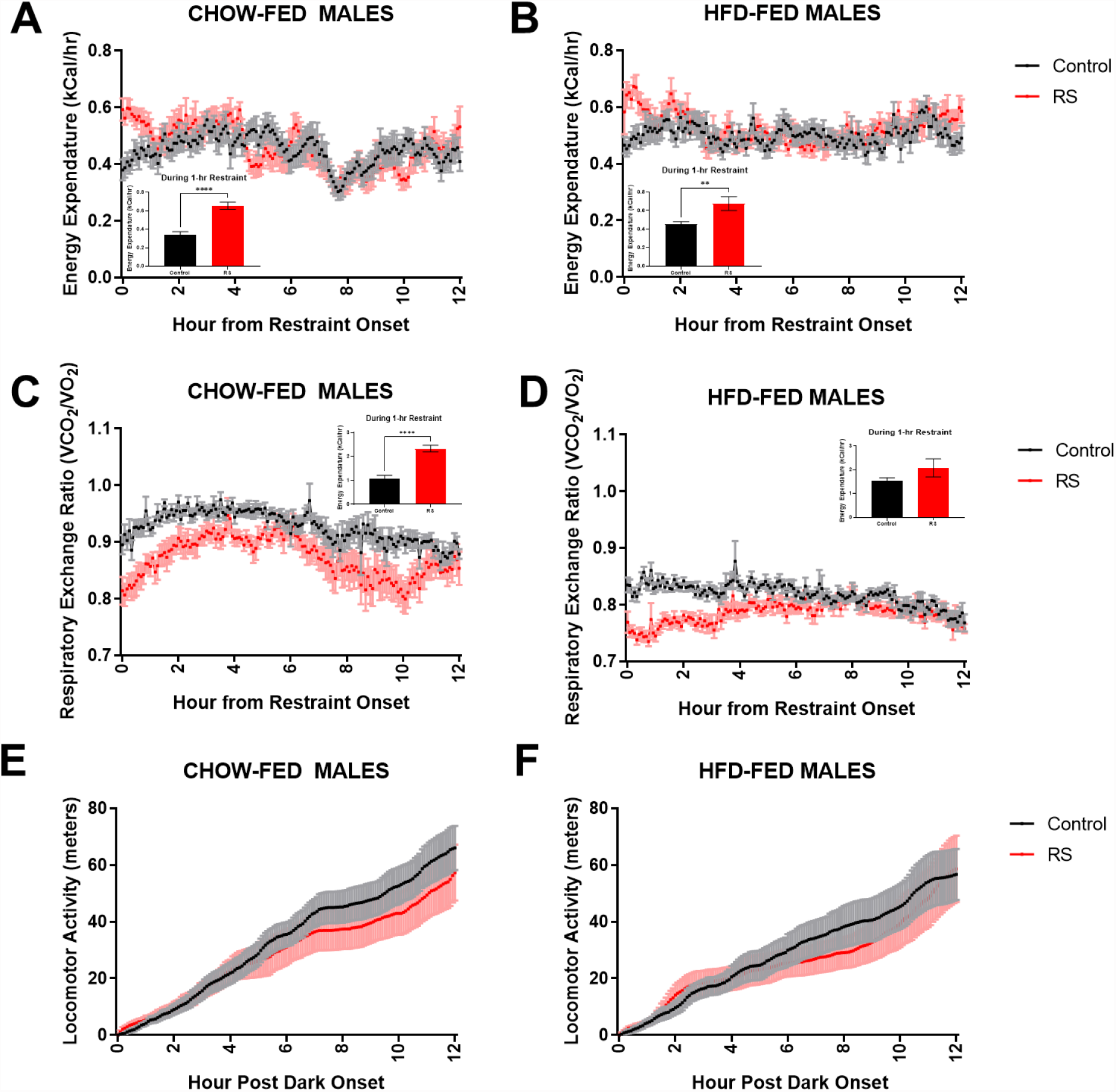
Metabolic Readings Following Restraint Stress or No Restraint Stress in Lean and Obese Male Mice. Energy expenditure (A, B), respiratory exchange ratio (C, D), and locomotor activity (E, F) of mice following no restraint (Control; black) or restraint stress (RS; red) which occurred for one hour prior to dark onset. Male wild-type mice were either chow-fed (A, C, E) or HFD-fed (B, D, F). **P < 0.01; ****P < 0.0001. N = 9-12/group.

Although stress reduced the respiratory exchange ratio (RER) in lean male mice (unpaired t-test, P < 0.0001; Figure 2C inset) during restraint, there was no significant effect of stress in lean mice during the 12h following the stress bout (Two-way ANOVA: F (144, 3190) = 0.7500, P = 0.9878, Figure 2C). However, there was a significant interaction of stress in obese mice during the restraint period and the 12h following stress (Two-way ANOVA: F (144, 2755) = 2.626, P < 0.0001, Figure 2D), though there was no significant difference during the one hour restraint (unpaired t-test, p = 0.1820; Figure 2D inset). This reduced RER is indicative of increased reliance on fat oxidation. Locomotor activity was not affected by acute restraint stress in either lean (Two-way ANOVA: F (216, 4774) = 0.1445, P > 0.9999; Figure 2) or obese mice (Two-way ANOVA: F (222, 4237) = 0.1692, P > 0.9999, Figure 2).

### 3.2 Activity of dLS Glp1r Neurons in Lean and Obese Mice During Restraint Stress

We used fiber photometry to assess real-time activity of dLS Glp1r-expressing neurons during acute restraint stress in lean and obese mice. Restraint stress rapidly stimulated the activity of Glp1r-expressing dLS neurons as shown by significantly increased peak area and spike frequency normalized to the pre-restraint stress period. This effect is easily observed in representative dLS Glp1r+ neuronal traces of one lean (Figure 3A and B) and one obese mouse (Figure 3C and D) during non-restraint and then during restraint the following day. Overall, obese male mice had a lower normalized spike frequency and normalized GCaMP calcium peak area (spikes only) from the GCaMP7 signal compared to lean male mice (Figure 4A and B). Two-way ANOVAs revealed a diet effect for both normalized spike frequencies (F [1,12] = 5.439, P = 0.038) and normalized peak areas (F [1,12] = 5.849, P = 0.0324; and Bonferroni’s multiple comparisons test revealed significant differences at segment two and three, P = 0.0364 and P = 0.011, respectively) in male mice. Contrasting this, the diet effect for normalized peak areas (F (1,20) = 0.661, P < 0.426) and spike frequencies (F [1,20] = 0.134, P < 0.718) was not significantly different between lean and obese female mice (Figure 4C and D).

**Figure 3.**
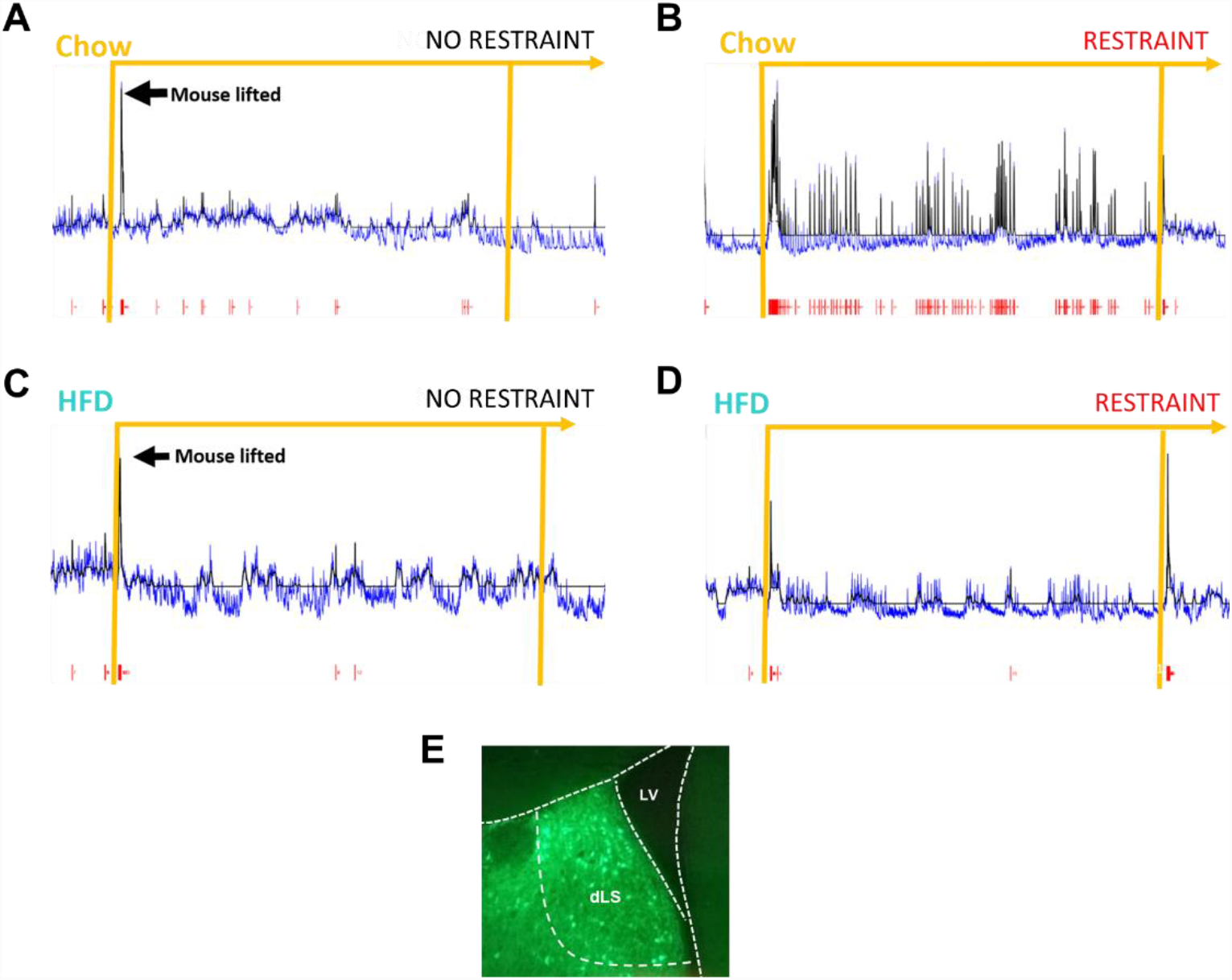
Representative Recordings of dLS Glp1r Neuronal Traces. Chow (A, B) and HFD-fed (C, D) fluorescent signal, a proxy for neuronal activity, during no restraint (A, C) and restraint (B, D). Orange line and initial spike in fluorescent signal occurred when mouse was lifted by the tail to simulate putting the mouse in a restrainer (NO RESTRAINT [A, C]) or to put the mouse in a restrainer (RESTRAINT B, D). The mLspike algorithm, which is a validated way to impute spike frequency based on Ca2+ transients, was used to measure spike (red hatch marks) frequency in MATLAB and generated the ‘best trajectory’ of calcium values (black lines) (Deneux et al., 2016). E. Representative photomicrograph of GCaMP expression in the dorsal lateral septum (dLS), 0.98mm from bregma. LV = lateral ventricle.

**Figure 4.**
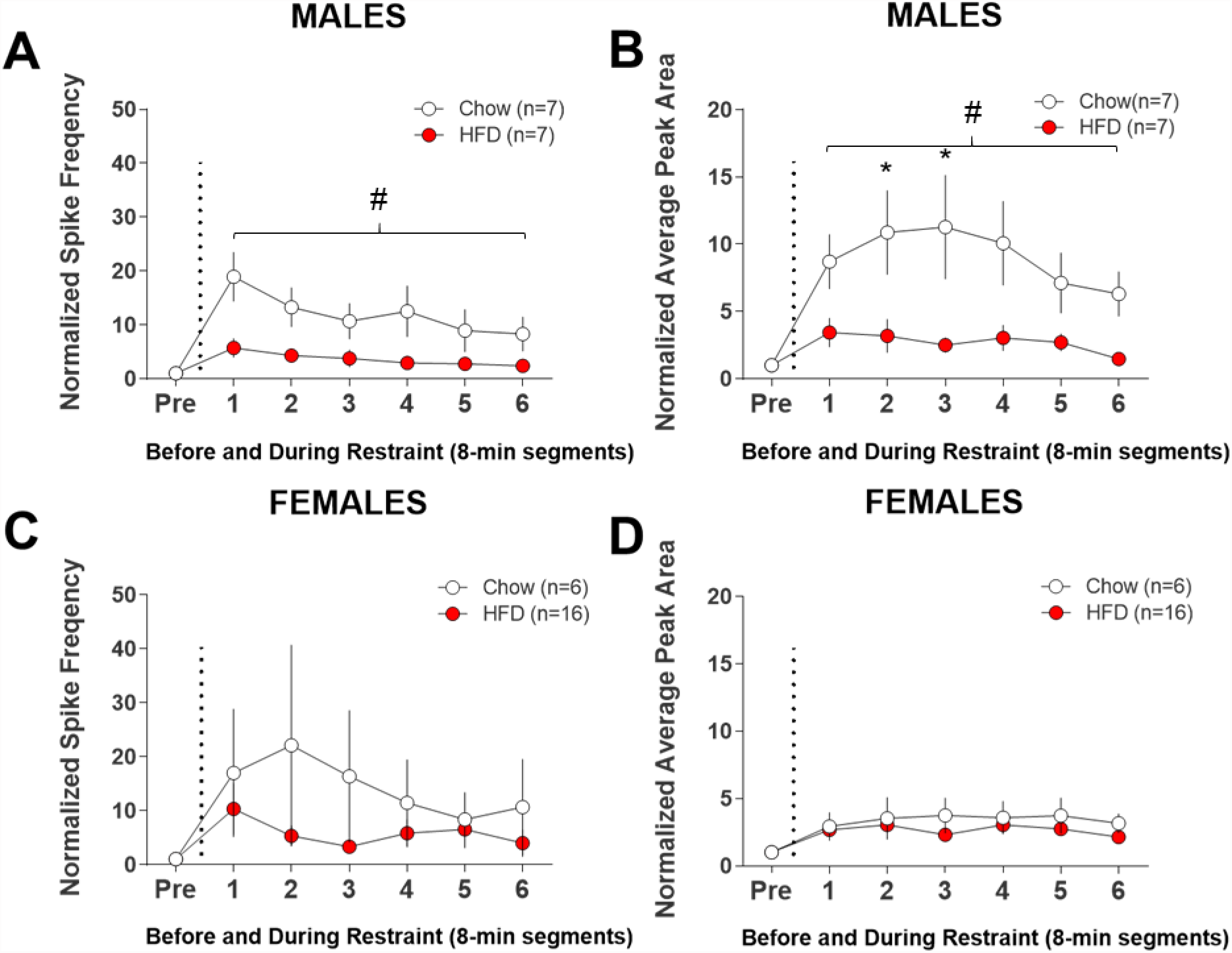
Restraint Stress Increases dLS Glp1r Neuronal Activity in Lean Male Mice but Not in Obese Male or Lean or Obese Female Mice. Fiber photometry recordings of Glp1r neurons in the dLS in male (A, B) and female (C, D) Glp1r-Cre mice. Recordings adjusted for baseline fluctuations and segmented into six bins to assess average frequency per minute of calcium spikes (A, C) and area under the peaks (B, D), in arbitrary units, normalized to pre-restraint. #P < 0.05 in a Two-way ANOVA; *P < 0.05 in a t-test. N = 6-16/group.

### 3.3 Head Movement Analysis

One factor that could explain reduced activation of Glp1r-expressing dLS neurons in obese mice during stress is that restraint may not be as physically stressful to obese mice as it is to lean mice. During restraint stress, rodents engage in periodic struggle behavior characterized by head and/or tail movement. This struggle behavior is believed to be representative of active escape behavior and is associated with increased neuronal activity in stress-related brain regions such as the BNST (Luchsinger et al., 2021). We used a custom restraint stress apparatus that enabled analysis of head movement during the fiber photometry experiments and used this as an index of the degree of stress experienced by lean and obese mice. Paired t-tests with Bonferroni correction revealed no difference in average length of struggle bouts (males, p = 0.966; females, p = 1.0; Figure 5A), number of head movement bouts (males and females, p = 1.0; Figure 5B), or total struggle time in a bout (males and females, p = 1.0; Figure 5C), between lean and obese mice. Taken together, the absence of any differences in struggle behavior between lean and obese male or female mice suggests that lean and obese mice experience restraint stress similarly.

**Figure 5.**
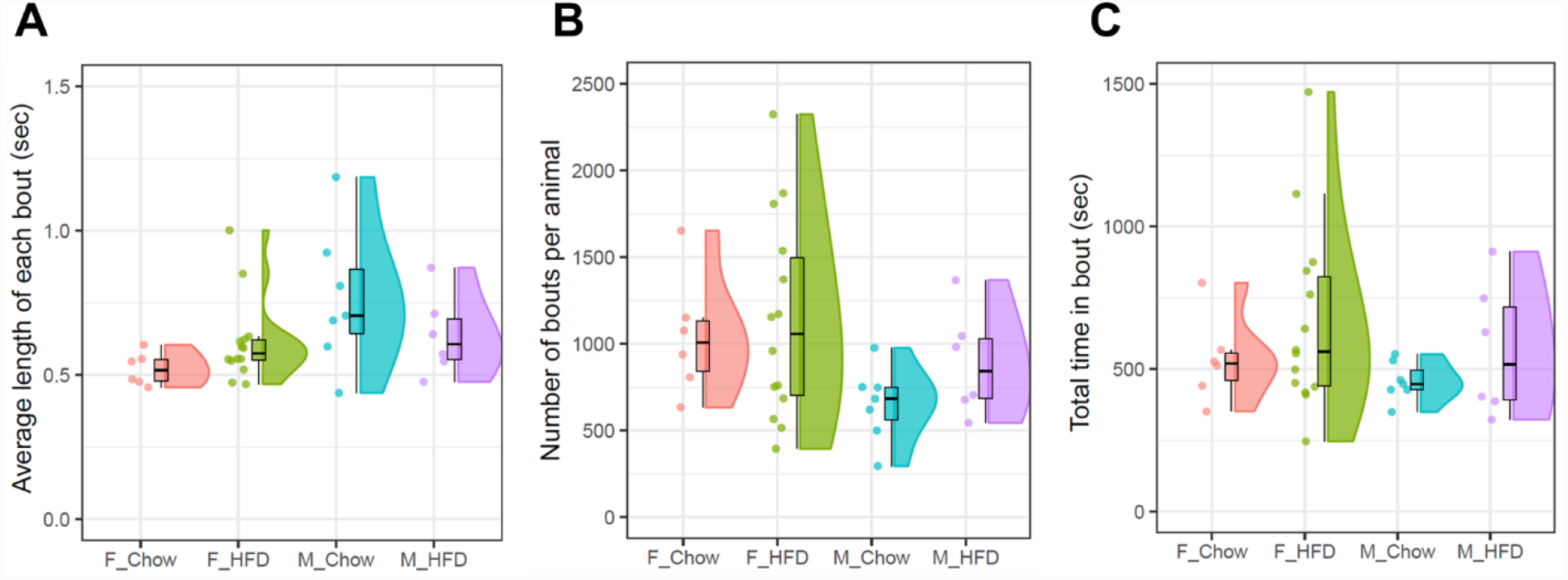
Struggle Behaviors Show Lean and Obese mice Struggled Similarly. Struggle analysis showed no significant differences between dietary conditions within sex suggesting that chow-fed and HFD-fed mice experienced the restraint stress similarly. Average length of each bout (A), number of bouts per animals (B), and total time in bout (C) were analyzed (N = 6-14/group).

Measuring head movement during the fiber photometry experiments allowed for direct comparison of time-locked active struggling to calcium transients measured in Glp1r-expressing dLS neurons (Figure 6A, B). In this analysis, time = 0 represents the onset of the struggle bout, and GCaMP activity is shown for the 5 seconds before and after the onset of the struggle bout. Comparing head movements time-locked with fiber photometry recording across maximum peaks show that calcium transient amplitude increases prior to the struggle onset in both lean and obese male mice, but lean males had a greater max peak than obese males after the initiation of the struggle bout (paired t-tests with Bonferroni correction, p = 0.007; Figure 6A). There was no difference in maximum peaks between lean and obese females (p = 1.0; Figure 6B). Analysis of area under the curve (AUC) shows greater AUC in lean vs. obese males (paired t-tests and Bonferroni correction p = 0.003; Figure 6C), but no difference between lean and obese females (p = 1.0; Figure 6C). Taken together, these data suggest that lean and obese males experience restraint stress equally (i.e. there is no difference in struggle behavior), but the response of dLS Glp1r neurons is increased in lean, male mice. Furthermore, there are no apparent differences between lean and obese female mice. Since dLS Glp1r signaling contributes to acute restraint stress-induced hypophagia in mice (Terrill et al., 2019), these results also align with our observation that stress-induced hypophagia is blunted by obesity in male but not female mice.

**Figure 6.**
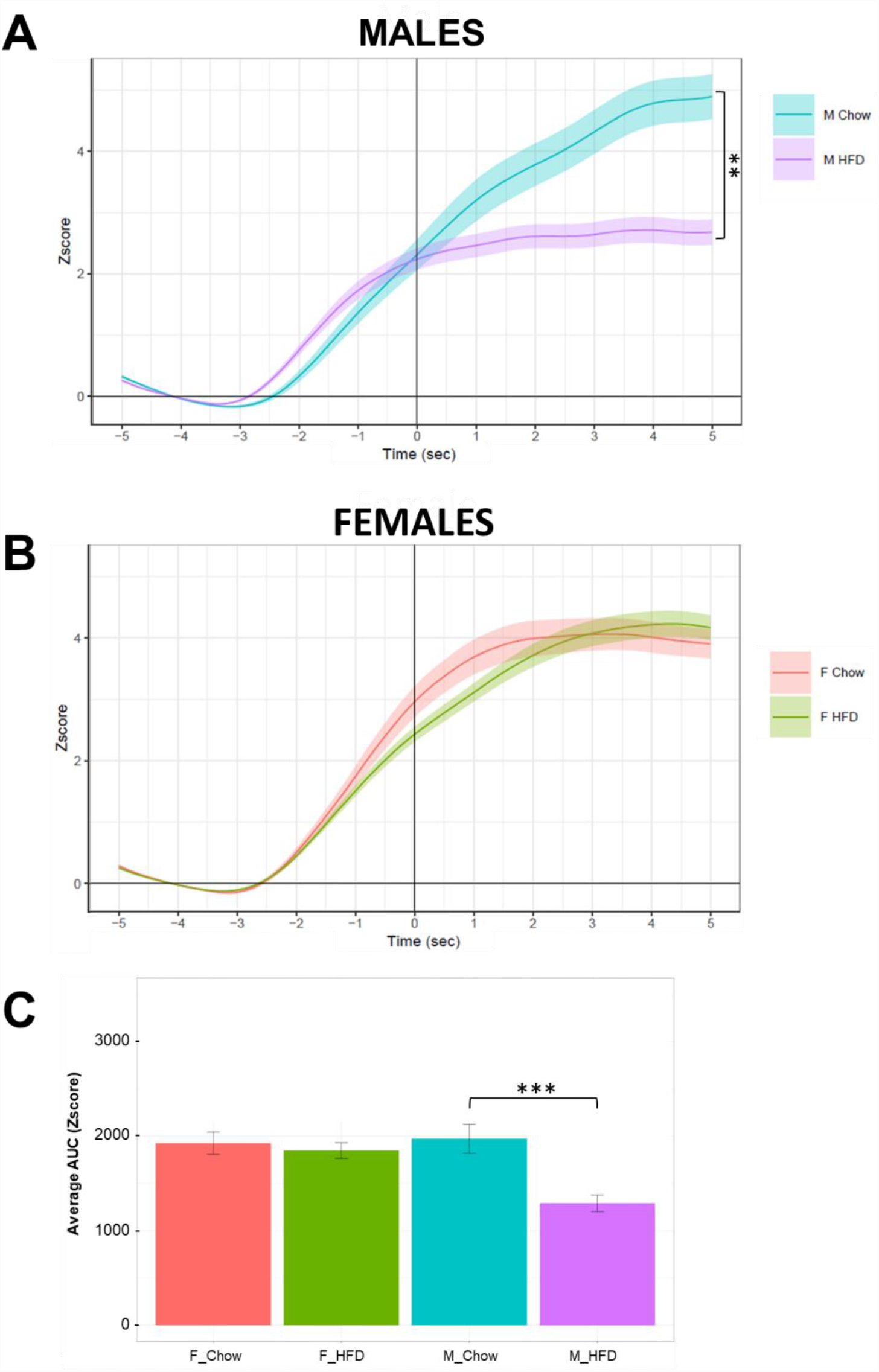
dLS Glp1r Neurons are Activated during Struggle in Lean Male Mice compared to Obese Male Mice, whereas Neuronal Responses in Lean and Obese Female Mice do Not Differ during Struggle. (A) Male and (B) female fiber photometry recordings time-locked to the struggle analysis. Maximum peaks were calculated for the 5 s after bout onset to normalize across bouts and time-locked to the fiber photometry recordings via MATLAB. Movement bout begins at zero seconds. Shaded area is SEM. (C) Average AUC by group. **P < 0.01; ***P < 0.001. (N = 6-14/group).

### 3.4 Effect on Food Intake Following Activation or Inhibition of dLS Glp1r Neurons

The observation that obese male mice display impaired stress-induced hypophagia and activation of LS Glp1r neurons does not demonstrate causation. To link the food intake responses to the dLS Glp1r neuronal response, we used chemogenetic modulation of LS Glp1r neurons. To determine whether activation of dLS Glp1r neurons is sufficient to reduce food intake, we tested whether chemogenetic activation of Glp1r-expressing neurons in the dLS mimics the effect of stress to induce hypophagia. Activating dLS Glp1r neurons with a G_q_-coupled DREADD did not significantly decrease food intake (Two-way ANOVA: F (11, 110) = 0.5334 ; P = 0.8769 ; Figure 7) in male mice. Male mice with control virus injected into the dLS had no difference in food intake in saline vs. CNO injected mice (Two-way ANOVA: F (1, 16) = 0.2511; P = 0.6231; not shown).

**Figure 7.**
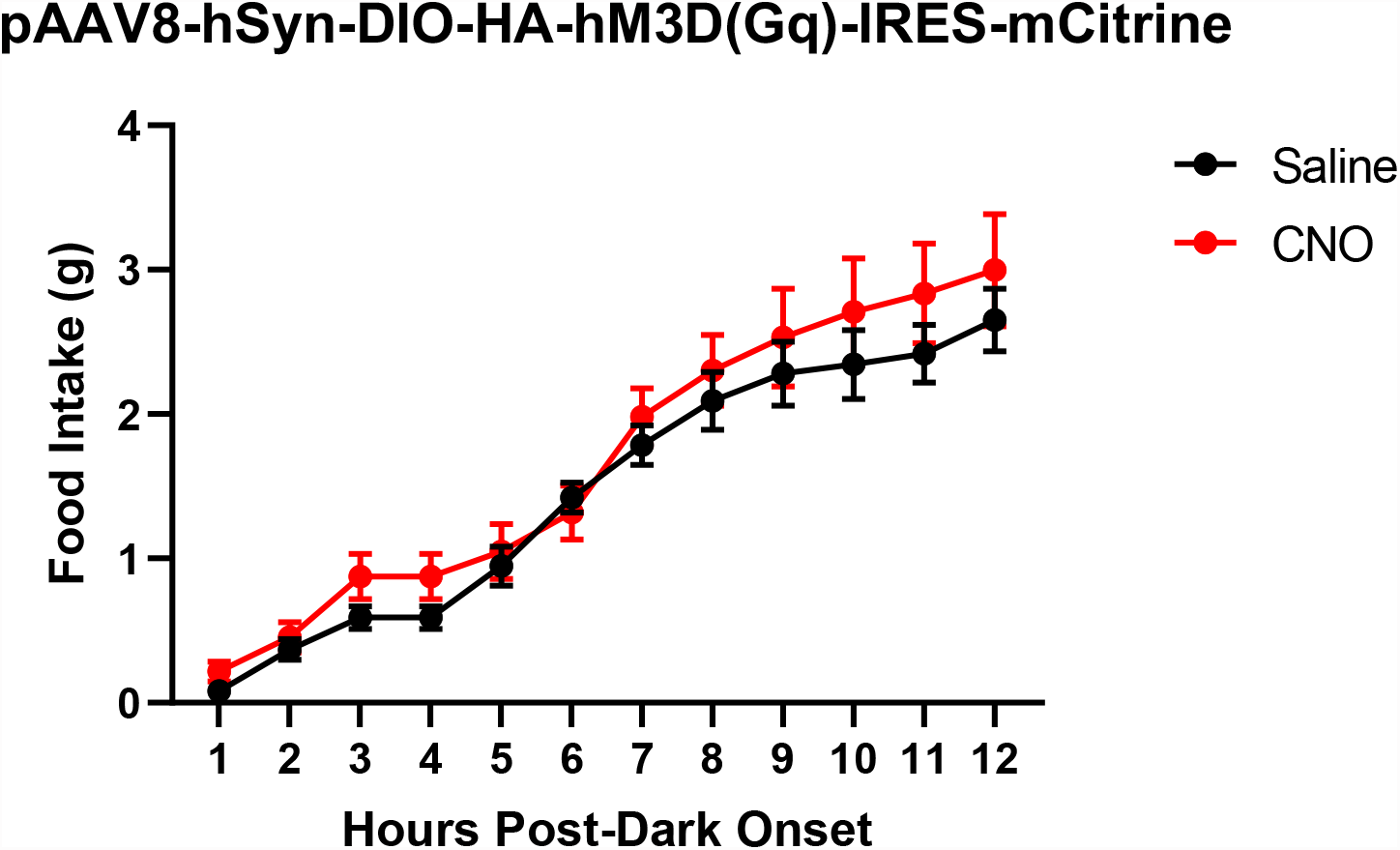
Activating Glp1r Neurons in the dLS does Not Reduce Food Intake. Food intake measurements in male mice with activating DREADD in the dLS. CNO (2 mg/kg) or saline injections (i.p.) were administered 30 min prior to restraint or no restraint for one hour before dark onset. (N = 6/group).

To determine whether activation of dLS Glp1r neurons is necessary for acute stress-induced hypophagia, we combined restraint stress with chemogenetic inhibition of dLS Glp1r neurons using a G _i_-coupled DREADD. Restraint stress promoted hypophagia in saline control mice, although under these conditions, the effect of restraint stress was not statistically significant (Two-way ANOVA: F (5, 60) = 1.402; P=0.2366; Figure 8A). Nevertheless, inhibiting dLS Glp1r neurons completely blocked stress-induced-hypophagia (Two-way ANOVA: F (5, 60) = 0.4834; P=0.7873; Figure 8B). These findings suggest that dLS Glp1r neurons are necessary for acute restraint stress-induced hypophagia.

**Figure 8.**
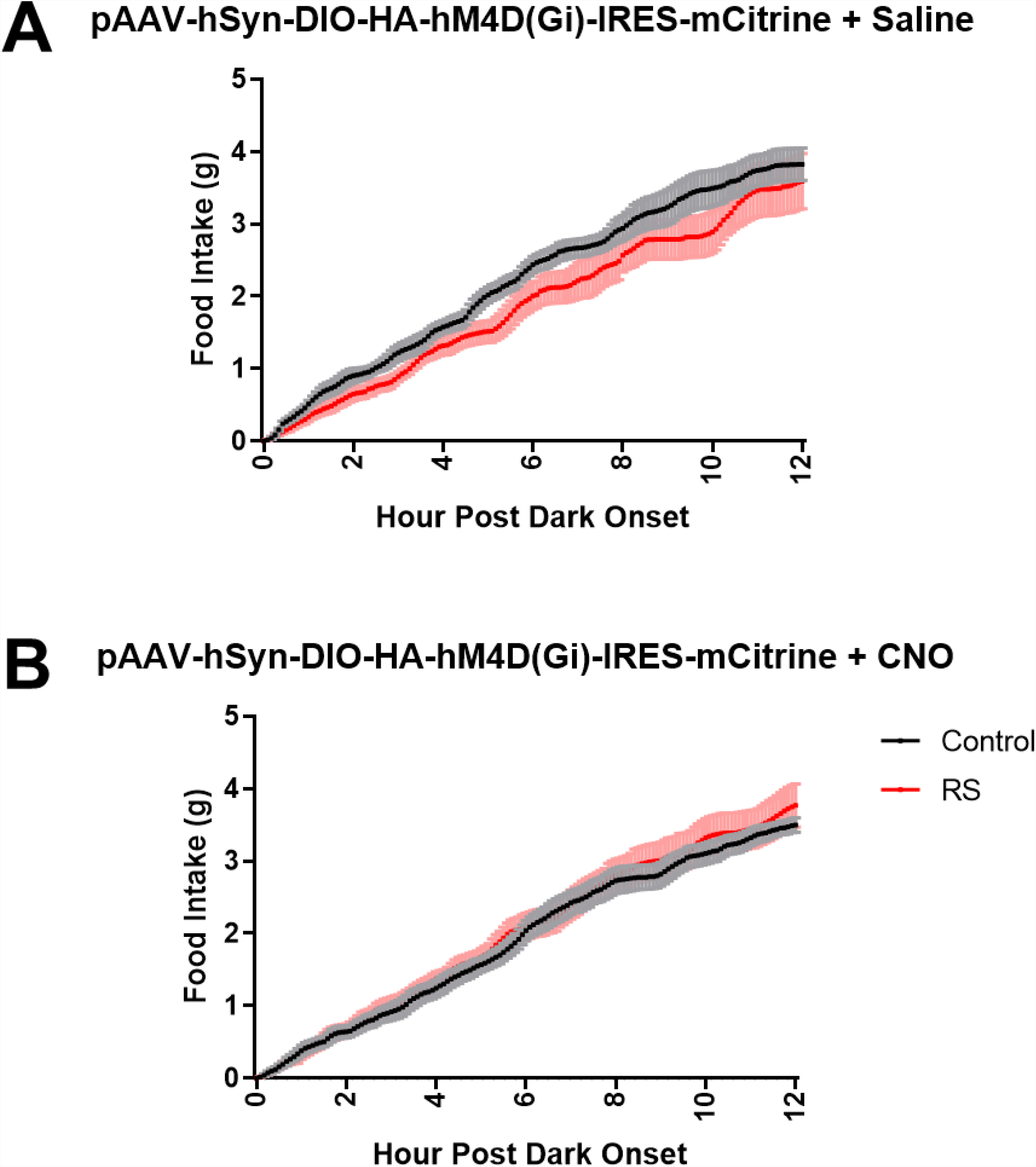
Inhibiting dLS Glp1r Neurons Blunts Stress-Induced Hypophagia. Food intake after male mice were injected with inhibitory DREADD into the dLS and following saline (A) or CNO injection (B; 2 mg/kg, i.p., bottom) 30 min before either restraint or no restraint which occurred one hour before dark onset. (N = 7-11/group).

## 4.0 Discussion

Although stress can promote weight gain and obesity, little is known about the effects of stress in the context of already established obesity. Clinical studies suggest that obese individuals are more susceptible to the negative effects of stress, such as depression, which can lead to a deleterious feed-forward cycle of “comfort feeding” (Slochower et al., 1981; Boggiano et al., 2015; Laitinen et al., 2002; Barrington et al., 2014, Coulthard et al., 2021). This provides an opportunity to identify mechanisms that regulate caloric intake in response to stress and how these mechanisms may be altered in the setting of obesity. We used a combination of sophisticated metabolic phenotyping and real-time neuronal activity measurements to show for the first time that Glp1r-expressing neurons in the dLS are rapidly and robustly activated during acute restraint stress in lean male mice, and this is markedly attenuated in obese male mice. Our subsequent observation that chemogenetic inhibition of dLS Glp1r neurons attenuates acute restraint stress-induced hypophagia suggests that impaired activation of these neurons in obesity contributes to increased caloric intake in response to stress. This identifies a novel target that can explain deleterious changes in feeding behavior at the intersection of stress and obesity.

Our results are in line with previous studies showing that inhibition of Glp1r signaling in the dLS attenuates restraint hypophagia in rodents (Terrill et al., 2018 & 2019). This suggests that acute restraint stress-induced hypophagia requires stimulation of Glp1r signaling and the activity of Glp1r-expressing neurons. One possibility is that stimulation of Glp1r signaling enhances the activity of dLS Glp1r-expressing neurons as has been previously shown in Glp1r-expressing neurons in the paraventricular hypothalamus (Liu et al., 2017). The LS is innervated by Glp1-producing NTS PPG neurons (Rinaman, 2010; Williams, 2018), so if obesity impairs Glp1 production or another function of NTS PPG neurons, this could lead to attenuated activation of dLS Glp1r neurons. Interestingly studies show that expression of the Glp1-producing PPG gene in the NTS is actually increased in obese rodents (Barerra et al., 2011; Vrang et al., 2007). This raises the possibility that obesity is associated with central Glp1 resistance, thus contributing to reduced activation of dLS Glp1r neurons.

Previous studies on the effect of obesogenic diets on stress and feeding behavior have primarily provided substances such as sucrose or HFD either shortly before or concomitant with the onset of stress. Short-term access to these substances generally impairs stress-induced hypophagia (Martin & Timofeeva, 2010; Michel et al., 2003; Moles et al., 2006; Pecoraro et al., 2004; Chuang et al., 2011; Bartolomucci et al., 2009). Novelty, palatability, and rewarding aspects of sucrose and fat are likely to be key contributors to increased intake even in the face of stress. It is not clear whether and to what degree these characteristics are still relevant following chronic exposure to HFD in the present studies. Thus, we cannot conclude that obesity itself rather than exposure to the HFD is what impaired stress-induced hypophagia in the present studies. Future experiments will test whether short-term exposure to HFD impairs stress-induced hypophagia in mice presented with either HFD or chow after restraint stress. Furthermore, it will be of interest to determine the effect of short-term exposure to HFD on stress-induced LS Glp1r neuronal activity. Restraint stress induced activation of LS neurons is attenuated in rats that had been previously provided intermittent access to sucrose (Martin & Timofeeva, 2010; Michel et al., 2003), raising the possibility that short-term exposure to palatable substances specifically mitigates the response of LS Glp1r-expressing neurons to stress.

Our observation that chemogenetic inhibition of dLS Glp1r neurons attenuates acute restraint stress-induced hypophagia suggests that activation of these neurons is necessary for the hypophagic effect of stress. This leads us to propose that stress activates dLS Glp1r neurons, consequently resulting in hypophagia, but in the setting of obesity, the impaired activation of dLS Glp1r neurons directly contributes to the attenuation of acute stress-induced hypophagia. This assigns a key role for dLS Glp1r neurons in the regulation of caloric intake following stress. It is, therefore, surprising that chemogenetic activation of dLS Glp1r neurons did not suppress caloric intake as we hypothesized. This contrasts a recent finding by Azevedo et al. (2020) showing that chemogenetic activation of LS Glp1r neurons reduces food intake. There are some key differences to highlight between our studies and those of Azevedo et al. (2020). First, expression of our activating DREADD was under the control of the human synapsin (hSyn) promoter expressed only in neurons, whereas Azevedo and colleagues used a DREADD construct driven by the elongation factor-1 alpha (Ef1a) active in multiple cell types. Second, we specifically targeted activating DREADDs to the dLS whereas Azevedo and colleagues (2020) targeted DREADDs to more ventral portions of the LS. This latter difference raises the possibility that neurons in different subdivisions of the LS exert specific effects on caloric intake.

Obesity elevates basal corticosterone levels in rodents, and stress subsequently increases corticosterone levels equally in lean and obese mice (Tannenbaum et al., 1997; Balsevich et al., 2014; Kalyani et al., 2016; Sharma and Fulton, 2013; Appiakannan et al., 2020). This suggests that lean and obese rodents experience stress equally. However, since we used larger restrainers to accommodate obese mice for our stress tests, a possible explanation for the blunted stress-induced hypophagia and dLS Glp1r neuronal activation in obese mice is that obese mice were simply not as stressed in the larger restrainers. We used two types of noninvasive measurements to assess the degree of stress experienced by all groups. Performing experiments in metabolic cages demonstrated that EE increases during the restraint stress bout. We interpret this increase in EE as resulting from struggling behavior (e.g. increased muscle contractions and breathing) while in the restrainer. The observation that EE increased equally in both lean and obese mice during restraint suggests that lean and obese mice equally struggle during this procedure. This was verified by our innovative video-based DLC analysis showing that there was no difference in struggle behavior between lean and obese mice. The ability to time-lock GCaMP7 signals to struggle behavior revealed an interesting response. For each struggle bout, dLS Glp1r neuronal activity increased equally between lean and obese mice beginning 2-3 seconds prior to struggle onset. Whereas dLS Glp1r neuronal activity continued to rise in lean mice after the onset of struggle, this neuronal activity plateaued upon struggle onset in obese mice. This suggests that activity of dLS Glp1r neurons is coupled to struggle behavior, and obesity interferes with this coupling.

One of our novel findings is that female mice are resistant to the hypophagic effects of stress regardless of whether they are lean or obese. The fact that there was also no difference in dLS Glp1r neuronal activity during stress between lean and obese female mice further supports the proposed role for dLS Glp1r neurons in the regulation of post-stress feeding. We cannot exclude the possibility that stress can affect feeding behavior and/or dLS Glp1r neuronal activity during specific phases of the estrous cycle. Egan and colleagues showed that a limited sucrose intake paradigm reduces the hypothalamic-pituitary-adrenal (HPA) axis response to stress during proestrus/estrus but not during diestrus 1/diestrus 2 (Egan et al., 2019). Including more female mice in the fiber photometry experiments and measuring estrous cycles or recording only on specific days of the cycle would help determine any differences in dLS Glp1r responses across the cycle and between dietary conditions. From an evolutionary perspective, one advantage of females maintaining normal food intake under stressful conditions could be to prepare for or to sustain a pregnancy. The LS also has other sexually dimorphic functions. Only female juvenile mice with vasopressin administered into the LS reduce social play behaviors, and male juveniles with vasopressin injected into the LS increase anxiety-like behaviors (Bredewold et al., 2014; see Bangasser et al., 2021 for review of other sex differences in mechanisms of anxiety).

Taken together, our results combine novel approaches to show that stress activates dLS Glp1r neuronal activity in lean male mice, but obesity is associated with impaired activation of dLS Glp1r neurons during stress and, as a consequence, attenuated hypophagia. In response to chronic stress, this could provide a mechanism for increased susceptibility to stress in obese individuals and further exacerbate weight gain. Future studies will focus on downstream targets of the LS that regulate feeding behavior. The LS extends projections to brain regions involved in the regulation of food intake, such as the lateral hypothalamus. Interestingly, LS Glp1r neurons are GABAergic (Risold and Swanson, 1997; Zhao et al., 2013), suggesting that stress-induced activation of LS Glp1r neurons exerts inhibitory control on post-synaptic targets. Although we focus on obesity in these studies, it is clear that obesogenic substances also impair both the hypophagic response to stress and activation of LS neurons in the absence of obesity (Martin & Timofeeva, 2010; Michel et al., 2003). Future studies will also determine whether exposure to obesogenic substances in the absence of obesity specifically attenuates stress-induced activation of LS Glp1r neurons and whether this contributes to impaired hypophagia following stress.

## Abbreviations

Glp1: Glucagon-like peptide-1
Glp1r: Glucagon-like peptide-1 receptor
dLS: dorsal Lateral Septum
PPG: preproglucagon
NTS: nucleus tractus solitarius

## Acknowledgements

The Vanderbilt Mouse Metabolic Phenotyping Center (MMPC), especially Louise Lantier, PhD and Merrygay James, helped run all experiments that used the Promethion (Sable Systems International). We also had assistance with the fiber photometry experiments from Gregory Salimando, PhD (baseline analysis), Andrew Gaulden, Amanda Morgan, PhD (AUC fiber photometry analysis), and Kellie Williford (restraint device design).

